# DIVERSITY in binding, regulation, and evolution revealed from high-throughput ChIP

**DOI:** 10.1101/122325

**Authors:** Sneha Mitra, Anushua Biswas, Leelavati Narlikar

**Affiliations:** Chemical Engineering Division, CSIR-National Chemical Laboratory, Dr. Homi Bhabha Road, Pune 411008, India

## Abstract

A high-throughput chromatin immunoprecipitation (ChIP) experiment is like a black-box: it reports all regions that are associated with the profiled protein based on the initial cross-linking step. These regions can be a highly diverse set of DNA sequences, with some making direct contact with the protein, some binding through intermediaries, and some being a result of long-range interactions involving the protein. We present diversity, a method that identifies the distinct components of such a mixture, leaving no data behind, while at the same time, using no prior motif knowledge. Using the example of the REST protein, we show that these different components give insights into the various complexes that may be forming along the chromatin and their regulatory functions.

http://diversity.ncl.res.in/ (webserver)

https://github.com/NarlikarLab/DIVERSITY (standalone for Mac OSX/Linux)

## 1 Introduction

DNA regulatory regions are known to play key roles in a wide range of biological processes (Maston et al., 2006) by interacting with specific proteins. To better understand these regions, high-throughput chromatin immunoprecipication (ChIP) technologies are extensively used to map protein-DNA interactions across different cell-types and organisms. However, in spite of these efforts, we still do not know how regulatory information is encoded in the four-letter alphabet of our genome (Shlyueva et al., 2014). We attribute this to the manner in which data from high-throughput experiments are currently interpreted: although evidence points towards multiple distinct regulatory mechanisms being at play at any given point in time (Struhl, 1991), a common characteristic is sought from the data. Motif finding is one such glaring example, where a common sequence signature, typically a position weight matrix (PWM) (Staden, 1984), is learned from protein-DNA binding data under the assumption that the solution must be “overrepresented” in the full set. However, a protein can exert its influence on the DNA in more than one way, by changing co-factors, or through intermediaries, at times never making direct DNA contact. It can therefore adopt different configurations at different DNA locations causing the dataset to be highly diverse (Fig. 1). Even motif discovery methods that identify multiple motifs do not account for this kind of diversity because they use the traditional enrichment criterion: the first “most enriched” motif in the complete set is masked before finding the second one, which is masked next and so on. The motifs “together” do not explain the dataset.

To address this issue, we developed an unsupervised learning approach that identifies different modes of protein-DNA binding from ChIP data (Narlikar, 2013). Here, we present diversity, which builds on that work, making it scalable to large datasets by reducing the number of models to be data partitioned based on four diverse binding modes learned. Through the example of RE-1 silencing transcription factor (REST), we show that the applicability of diversity goes beyond merely identifying the direct DNA-binding specificity of a protein: it can learn potentially diverse regulatory mechanisms supported by distinct evolutionary and functional characteristics.

**Figure 1:**
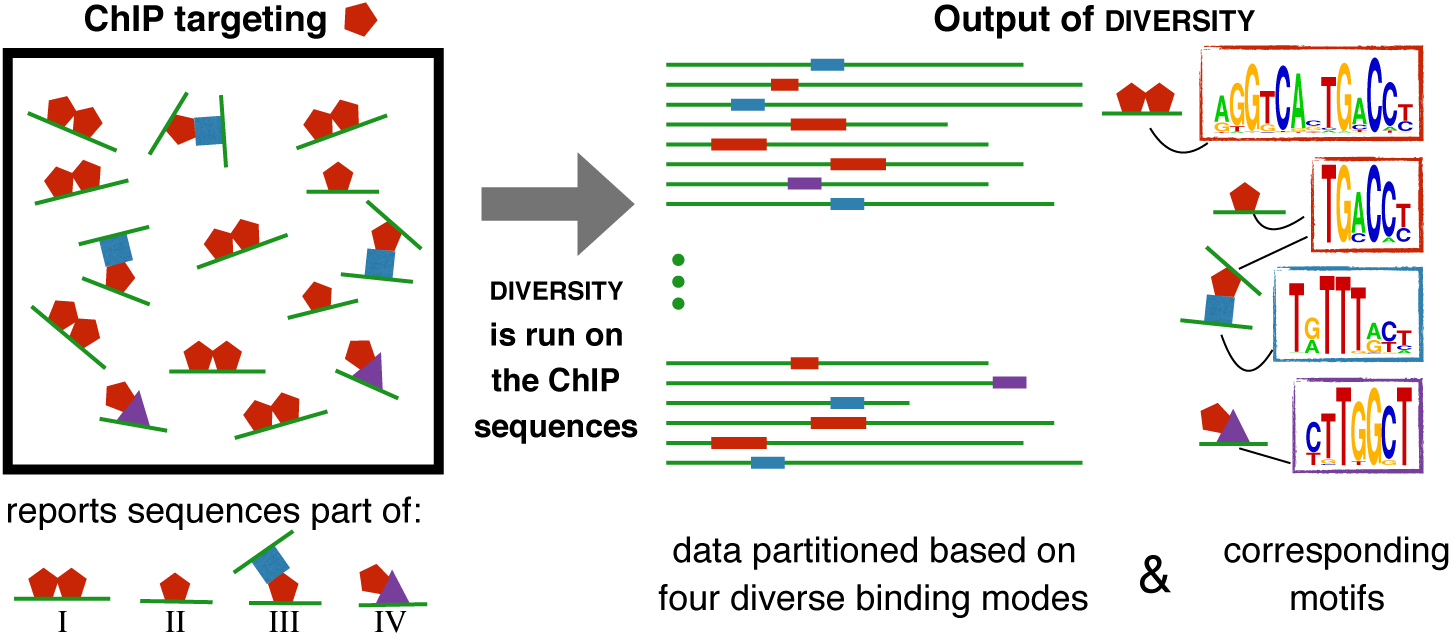
ChIP pulls down all complexes involving the profiled protein. Diversity splits reported regions into different sets based on motifs common to each set, all learned de novo. In this toy example, the red protein binds to DNA as (I) homodimer, (II) monomer, (III) indirectly via the blue protein (possibly causing loops), or (IV) indirectly to via the purple protein. The expected output of diversity is four modes.

## 2 Methods

### 2.1 Overview of Diversity

Diversity assumes the protein in the ChIP experiment makes m types of DNA-contacts, or has *m* DNA-binding modes, each represented as a PWM ((Fig. 1). It then partitions the dataset into *m* subsets, while simultaneously learning one enriched PWM corresponding in each subset. Crucially, the user needs to input nothing but the ChIP sequences. Everything else from the width of the motifs to the optimal number of modes is learned directly from the data using a mix of Gibbs sampling and Bayesian model selection.

Diversity makes use of multiple processing cores, which are now standard in most computers. This and other algorithmic advances detailed in Supplementary methods make diversity comparable in speed to the standard motif discovery tool MEME (Bailey et al., 2009) (Supplementary figure S2). However, we stress that the problem solved by diversity is different from any available tool: the goal is not to find individual motifs enriched in the set, but to explain the complete set using an optimal number of binding modes. Therefore, we cannot compare the output of diversity with these methods. Nevertheless, even if the goal is only to identify the direct binding site, we have previously established that diversity’s approach finds the direct binding motif of a protein if it makes DNA contact, as one of the modes; often when popular motif discovery methods fail (Narlikar, 2013). But it is the additional insights on looping, indirect binding, and regulatory function that make diversity a unique tool for downstream ChIP analysis.

### 2.2 Input and output of Diversity

The only mandatory input to diversity is the set of ChIP-reported sequences. A successful run of diversity produces an output directory containing the best model as an html file. Details of all learned models are stored as separate subdirectories, in case the user is interested in models with less (or more) binding modes. If input is given in a .bed format on the webserver, diversity reports additional information: multiple alignments of the sequences contributing to each mode, phastCons sequence conservation (Karolchik et al., 2014), and the distance of the modes to the closest TSS as box-plots (insights in (Fig. 1).

## 3 Diversity applied to REST

REST represses neuronal genes in non-neuronal cell-types and plays a role in differentiation of neuronal cells (Qureshi et al., 2010). It binds directly to a 21bp RE-1 motif, but is believed to interact with a diverse set of co-factors resulting in distinct transcription outcomes (Greenway et al., 2007). Diversity was run on REST data in neurons as compiled by Rockowitz et al. (2014), finding 12 modes ((Fig. 2). Rockowitz et al. applied MAST (Bailey et al., 2009), which scans sequences on the basis of a user-supplied PWM, to identify neuronal regions containing the RE-1 motif. They showed there was only a marginal enrichment of RE-1, even in the top 600 sequences. Therefore, it is not surprising that diversity also finds only a small fraction of sequences (< 4%) contributing to a mode (mode 1, (Fig. 2) that resembles RE-1. In addition to RE-1, diversity finds 11 other modes. For all these modes, genes with transcription start sites within 2kb are significantly highly expressed. This supports behaviour of REST as an activator in neurons (Ooi and Wood, 2007), but suggests this likely happens not by binding DNA directly, but via co-factors. Furthermore, modes differ in terms of distance from the closest gene, expression of the closest gene, as well as evolutionary characteristics. Co-factors potentially binding to the modes 2-12 are described in greater detail in the Supplementary information. Diversity was also run on the non-neuronal GM12878 cell line, where it finds only variants of the RE-1 motif. Sequence conservation and nucleosome occupancy, both indicate distinct functions for specific RE-1 variants over others (Supplementary figure S1).

**Figure 2:**
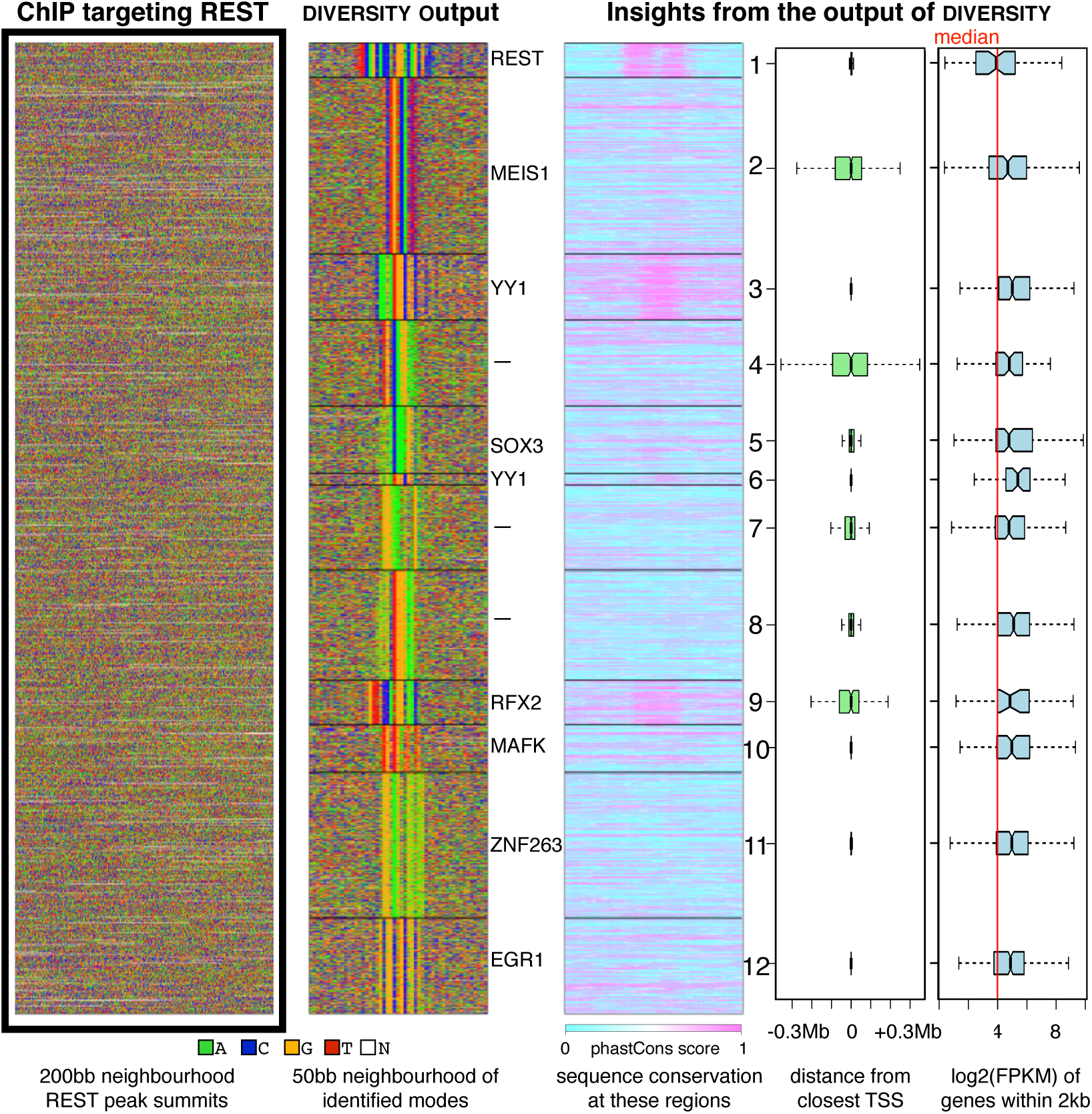
200bp centered around ChIP summits of REST are input to diversity, which finds 12 modes. Protein most likely to bind each mode (Gupta et al., 2007) is listed on the side. Corresponding sequence conservation, relative position, and gene-expression yields insights about the function of each mode.

## 4 Conclusion

Diversity asks the question: *what sequence component caused a specific region to be reported in a ChIP experiment?* The answer, in combination with additional data such as sequence conservation, SNPs, chromatin structure, downstream gene-expression, etc. can yield insights into the diverse regulatory mechanisms at play. The added benefits of a web-server and a standalone parallel version make diversity a practical tool for discovering new biology from ChIP experiments.

## Acknowledgements

We thank Rahul Siddharthan for useful discussions on the algorithm and the NCL systems administration team of Kishore Deshpande, Javed Rohile, and Avinash Deotale for their help in setting up the webserver.

## References

Bailey, T.L. et al. (2009) MEME SUITE: tools for motif discovery and searching. Nucleic Acids Res., 37, W202–208.

Greenway, D.J. et al. (2007) RE1 Silencing transcription factor maintains a repressive chromatin environment in embryonic hippocampal neural stem cells. Stem Cells, 25, 354–363.

Gupta, S. et al. (2007) Quantifying similarity between motifs. Genome Biol., 8, R24.

Karolchik, D. et al. (2014) The UCSC Genome Browser database: 2014 update. Nucleic Acids Res., 42, D764–770.

Maston, G.A. et al. (2006) Transcriptional regulatory elements in the human genome. Annu. Rev. Genomics Hum.Genet., 7, 29–59.

Narlikar, L. (2013) MuMoD: a Bayesian approach to detect multiple modes of protein-DNA binding from genome-wide ChIP data. Nucleic Acids Res., 41, 21–32.

Ooi, L. and Wood, I.C. (2007) Chromatin crosstalk in development and disease: lessons from REST. Nat. Rev. Genet., 8, 544–554.

Qureshi, I.A. et al. (2010) REST and CoREST are transcriptional and epigenetic regulators of seminal neural fate decisions. Cell Cycle, 9, 4477–4486.

Rockowitz, S. et al. (2014) Comparison of REST cistromes across human cell types reveals common and context-specific functions. PLoS Comput. Biol., 10, e1003671.

Shlyueva, D. et al. (2014) Transcriptional enhancers: from properties to genome-wide predictions. Nat. Rev. Genet., 15, 272–286.

Staden, R. (1984) Computer methods to locate signals in nucleic acid sequences. Nucleic Acids Res., 12, 505–519.

Struhl, K. (1991) Mechanisms for diversity in gene expression patterns. Neuron, 7, 177–181.

